# Clove-derived eugenol induces strong avoidance behaviour in the invasive fruit fly, *Drosophila suzukii*

**DOI:** 10.1101/2025.10.24.684488

**Authors:** Steve B. S. Baleba, Victor O. Omondi, Samira A. Mohamed, Merid N. Getahun, Nan-ji Jiang, Souleymane Diallo

**Author notes:** Corresponding Author’s.

## Abstract

The success of invasive species often relies on sensory adaptations that allow them to exploit new environments. Unlike most of its relatives, the spotted-wing drosophila (*Drosophila suzukii*) lays eggs in ripening fruit, making it a serious and rapidly spreading agricultural pest worldwide. Spice plants produce volatiles that repel many insects, making them a potential source of *D. suzukii* behavioural modulators. Here, we examined whether volatiles from clove (*Syzygium aromaticum*) could alter the behaviour and fitness of *D. suzukii*. Behavioural assays revealed strong aversion: adults avoided grapes treated with clove extract, while gravid females laid significantly fewer eggs on treated fruit in both no-choice and choice assays. Clove exposure adversely impacted fitness, resulting in slower larval growth, reduced adult weight, decreased emergence rate, and shortened lifespan. Gas Chromatography-mass spectrometry analyses identified eugenol as the predominant volatile. Consistent with this, synthetic eugenol elicited strong avoidance not only in grapes but also across various host fruits, including apple, banana, blueberry, guava, mango, orange, papaya, raspberry, strawberry, and tomato. Furthermore, molecular docking and single-sensillum recordings indicated that eugenol binds most strongly to DsuzOR88a, followed by DsuzOR69a and DsuzOR23a odorant receptors. Subsequently, cheminformatic screening of 1,000 eugenol analogues revealed several structural variants predicted to interact strongly with OR88a, offering additional candidate repellents. By combining behavioural, analytical chemistry, electrophysiological, and cheminformatic approaches, our study demonstrates that clove, through its major volatile eugenol, disrupts host use and reduces fitness in *D. suzukii*. These results highlight the ecological role of plant-derived repellents in insect–plant interactions and position eugenol as a promising natural compound for sustainable fruit pest management.

## Introduction

The spotted-wing drosophila (*Drosophila suzukii* Matsumura; Diptera: Drosophilidae) is an invasive pest of soft- and stone-fruit crops such as blueberry, strawberry, cherry, and grapes (Cai et al., 2019). Unlike most drosophilids that exploit decaying fruits, *D. suzukii* females possess a serrated ovipositor that enables oviposition in intact ripening fruits (Atallah et al., 2014; Galagovsky et al., 2024). This adaptation causes direct yield loss through larval feeding and further exposes fruits to secondary infestation by other drosophilids and opportunistic pathogens (Rombaut et al., 2017).

Originally described in Japan in 1916 (Kanzawa, 1939), *D. suzukii* has since spread worldwide, establishing itself in North America (Rota-Stabelli et al., 2020), Europe (Calabria et al., 2012; Ørsted & Ørsted, 2019), and Africa (Boughdad et al., 2020; Hassani et al., 2020). Multiple independent invasion events have been traced back to source populations in Asia, highlighting its repeated success in colonising new regions (Bolda et al., 2010; Goodhue et al., 2011). Recent surveys confirmed its establishment in East Africa (Kwadha et al., 2021), emphasising its ongoing global expansion. The broad ecological range of *D. suzukii* raises important questions about how behavioural and sensory adaptations contribute to invasion dynamics and host exploitation across new environments.

The economic impact of *D. suzukii* is severe. In the United States, early assessments estimated annual revenue losses exceeding $500 million in the West Coast berry and cherry industries (Bolda et al., 2010). Subsequent studies have quantified losses in diverse regions and examined the costs of prevention and control, including integrated pest management (IPM) compared to calendar-based insecticide programs (Walsh et al., 2011; De Ros et al., 2015; Farnsworth et al., 2017; DiGiacomo et al., 2019; Yeh et al., 2020; Knapp et al., 2021). Collectively, these studies demonstrate that *D. suzukii* imposes substantial yield losses and high management costs. Current management depends largely on insecticide sprays, but resistance development, regulatory restrictions, and negative environmental impacts challenge their long-term use (Asplen et al., 2015; Hamby et al., 2016). This underscores the need for alternative, eco-friendly strategies such as natural repellents derived from plant metabolites. Among potential natural sources, spice plants emit volatile compounds, especially phenolics, terpenes, and essential oils, that modify insect behaviour and performance (Fuentes-Lopez et al., 2025). These compounds represent an underexploited resource for pest management. Also, the sensory mechanisms governing their detection are poorly understood. For example, clove (*Syzygium aromaticum*) extract has been reported to repel or impair survival in diverse insect taxa, including psyllids and mosquitoes (Czarnobai De Jorge et al., 2022; Sanga et al., 2023; Lopez et al., 2025). However, in the context of *D. suzukii*, the effect of the specific active constituents on their behaviour and the olfactory basis of their detection remain unknown.

Odorant receptors (ORs) play a central role in translating chemical cues from the environment into neural signals that guide key behaviours such as host selection, oviposition, and avoidance (Yan et al., 2020). Here, we hypothesise that clove produces specific volatiles that repel *D. suzukii* and deter oviposition through activation of specific odorant receptors (ORs). We (1) quantified behavioural responses of adults and gravid females to clove extracts, (2) characterised volatile composition using gas chromatography–mass spectrometry (GC–MS), (3) assessed responses to individual clove-derived volatiles, (4) focused on eugenol as a dominant volatile and examined receptor–ligand interactions through molecular docking and single-sensillum electrophysiology, and (5) used cheminformatics to screen eugenol analogue volatiles as candidate repellents acting through specific ORs.

By integrating behavioural, analytical chemistry, electrophysiological, and computational approaches, our study examines how plant-derived volatiles interact with insect olfactory systems to shape host-use behaviour. This work demonstrates that clove volatiles strongly deter *D. suzukii* behaviour by binding to key odorant receptors and disrupting host selection, positioning it as a promising natural repellent for sustainable fruit pest management. It also highlights the uses of *D. suzukii* as a model to explore the broader ecological role of chemical cues in mediating insect– plant interactions and invasion processes.

## Materials and Methods

### Insect Rearing

We maintained a laboratory colony of *Drosophila suzukii* on a standard cornmeal–yeast diet at 25 ± 1 °C, 65 ± 5% RH, and a 12:12 h light: dark cycle. We used 3–5-day-old flies for all behavioural and electrophysiological assays. Gravid females were separated 24 h before the oviposition experiments.

### Preparation of Clove Solutions

We purchased clove (*Syzygium aromaticum*) powder from a local supermarket and used it to prepare aqueous suspensions at dilutions of 10^− 1^, 10^− 2^, 10^− 3^, and 10^−4^(w/v) by dissolving the powder in distilled water (Fig. 1a), followed by 15 minutes of sonication. We used distilled water as a control.

**Figure 1.**
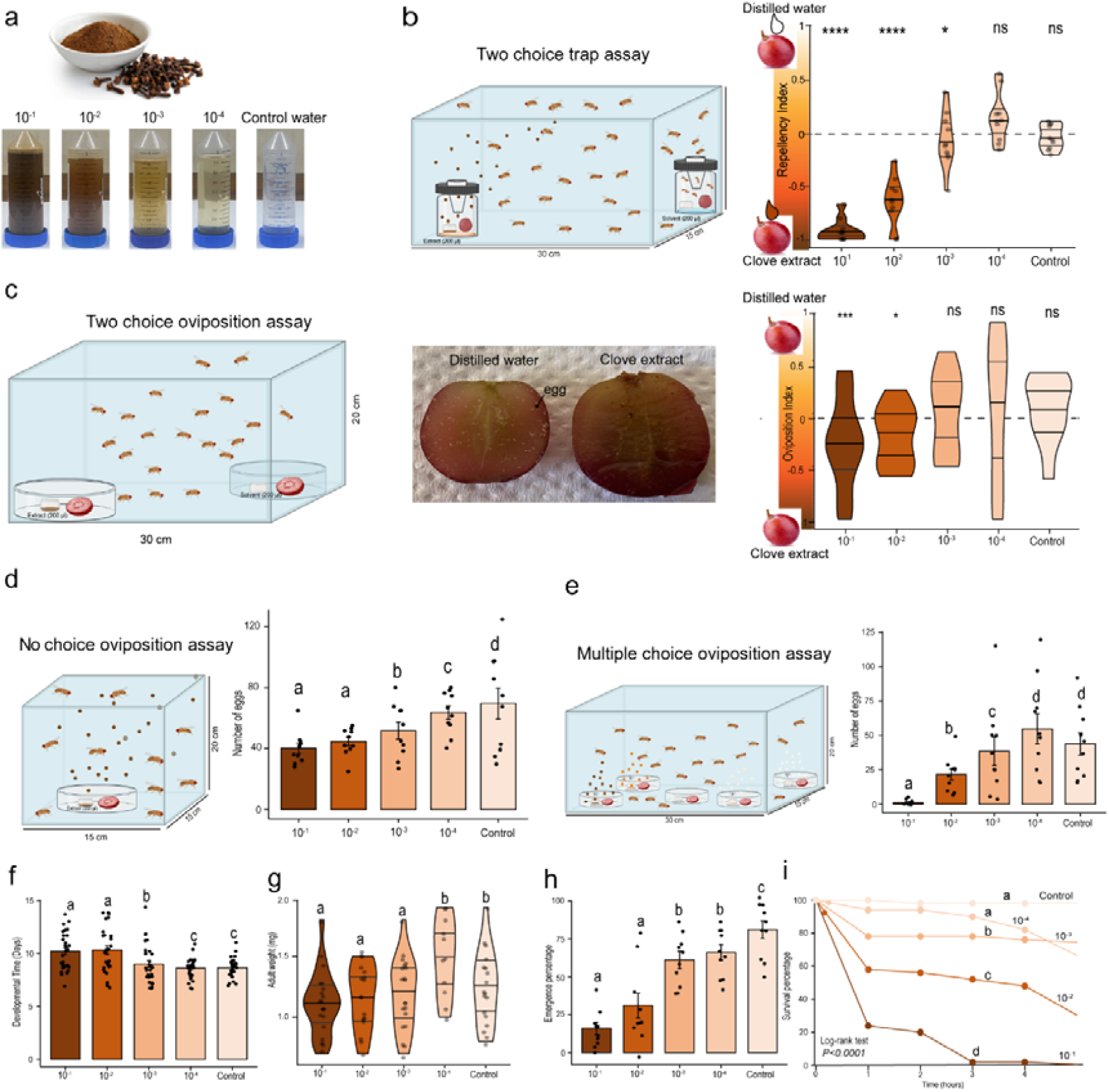
Clove extracts repel *Drosophila suzukii* adults and impair the performance of their offspring. **(a)** Photos illustrating how clove powder was serially diluted in distilled water. **(b)** Experimental setup for two-choice trap assays with grapes coated in clove extracts at different concentrations (10^− 1^–10^−4^). **(c)** Left panel showed the experiment sets. Photos showing reduced egg laying by females on grapes treated with clove extracts (right) compared to untreated grapes (left). Two-choice oviposition assay results showing reduced egg laying on clove-treated grapes as clove concentration increased. **(d and e)** No-choice and multiple-choice assay results showing reduced egg laying on clove-treated grapes with increasing clove concentration. **(f–i)** Life-history measurements showing slower larval development (f), reduced adult weight (g), lower emergence percentage (h), and shorter adult lifespan (i) in *D. suzukii* offspring developing on clove-treated grapes. Dots on each violin plot and bar graph represent data points from individual replicates. Error bars indicate the standard error of the mean (SEM). The symbol “n.s.” denotes a non-significant difference of the repellency index from the theoretical mean (0), while *, **, ***, and **** indicate significant differences with P□<□0.05, P□<□0.01, P□<□0.001, and P□<□0.0001, respectively. Bars or lines labelled with different letters differ significantly from each other.

### Two-Choice Trap Assay

We evaluated the repellency of clove solutions using two-choice trap assays (Fig. 1b). Groups of 30 mixed-sex adult flies (3–5 days old, starved for 24 h, ∼1:1 sex ratio) were gently released using a mouth aspirator into cages (15 cm × 15 cm × 20 cm) containing two traps. Each trap consisted of a 15 mL Nalgene vial (entry holes 5 mm diameter) baited with a half grape. One-half was coated with 200 µL of clove extract at a given dilution, while the control half was coated with distilled water. Trap positions were randomised between replicates to minimise side bias. Assays were conducted under controlled environmental conditions (25 ± 1 °C, 65 ± 5% RH, 12:12 h L:D). After 24 h, the number of flies captured in each trap was recorded. Each dilution (10^− 1^– 10^−4^ w/v) was tested with 10 independent replicates, each using new flies, fresh bait, and clean cages. Repellency was quantified using a preference index (PI = (T − C) / (T + C)), where T and C represent the number of flies in the treatment and control traps, respectively. The PI ranged from -1 (maximum avoidance) to 1 (maximum attraction). Zero thus denoted no choice.

### Oviposition Assays

We conducted no-choice, two-choice, and multiple-choice oviposition assays to evaluate the effects of clove solutions on *D. suzukii* egg-laying behaviour. In no-choice assays, a single half grape was treated with a clove extract (10^− 1^–10^−4^ w/v) or distilled water (control) and placed in a Nalgene vial without a lid within a 15 cm × 15 cm × 20 cm cage. Twenty gravid females were gently introduced using a mouth aspirator and allowed to lay eggs for 24 h, after which eggs were counted. In two-choice assays, one treated half-grape and one control half-grape were placed in separate Nalgene vials without lids and presented simultaneously in the same arena. Oviposition preference was quantified using the oviposition preference index (OPI = (T − C) / (T + C)), where T and C represent the number of eggs on the treated and control fruits, respectively. The OPI ranged from -1 (maximum avoidance) to 1 (maximum attraction). Zero thus denoted no choice. In multiple-choice assays, five half grapes, each coated with a different clove dilution or water, were offered simultaneously. Eggs on each fruit were counted after 24 h under a Leica S8 APO stereo microscope (Leica Microsystems, Germany) at 10–30× magnification. All assays were replicated 10 times using fresh flies, fruits, and clean cages for each replicate.

### Life History Trait Measurements

We assessed the effects of clove exposure on the developmental traits and adult survival of *Drosophila suzukii*. Eggs were obtained from laboratory colonies maintained on a standard cornmeal diet. Ten freshly laid eggs (0–3□h post-oviposition) were carefully collected using a fine brush and placed on each half grape treated with 200 µL of a specific clove solution (10^− 1^– 10^−4^ w/v) or distilled water (control). Treated grapes were placed individually in ventilated 50 mL plastic containers, covered with mesh and sealed with a rubber band, and maintained under controlled conditions (25 ± 1□°C, 65 ± 5% RH, 12:12 h L:D). Each treatment included 10 independent replicates, which were randomised across time and space to minimise positional or temporal bias. Larvae were allowed to develop on the grapes until adult emergence. We recorded developmental time from egg to adult, emergence percentage, and adult weight at eclosion, measured with a Sartorius analytical balance (±0.01 mg). To evaluate the effects of clove exposure on adult survival, three-day-old adults emerging from a standard diet were transferred in groups of 20 (1:1 sex ratio) into vials containing cotton soaked with the corresponding clove solution or distilled water (control). Mortality was recorded hourly until the death of all the adults.

### Chemical Analysis of Clove Extracts

We extracted volatiles from clove powder using dichloromethane (DCM) (Fig. 2a). One gram of powdered cloves, purchased from local supermarkets, was weighed in triplicate into 50 mL conical flasks. Each sample was mixed with 10 mL DCM and shaken for 2 h at room temperature on a mechanical shaker. The mixtures were filtered through 125 mm Whatman™ grade 1 filter paper, and the filtrates were subjected to GC–MS analysis. We injected 1 µL of each clove extract into a GC–MS system (GC HP-7890A, MS 597C, Agilent Technologies, USA) equipped with an autosampler (7683B series). Injections were performed in splitless mode, using helium (99.99% purity) as the carrier gas at a flow rate of 1.2 mL min^− 1^. Separation was achieved on an HP-5MS capillary column (30 m × 0.25 mm i.d., 0.25 µm film thickness, J & W Scientific, USA). The oven program was set to 35 °C (5 min hold), increased at 10 °C min^− 1^ to 280 °C, and held for 10.5 min. The ion source was maintained at 230 °C, with electron ionisation at 70 eV. Mass spectra were acquired over m/z 38–350 with a scan rate of 0.73 scans s^− 1^. Data were processed using Agilent MSD Productivity ChemStation software, with probability-based matching (initial peak width, 0.034; threshold, 15.7). Compounds were identified by comparing their retention times and mass spectra with those of the NIST 05a and Adams MS HP libraries.

**Figure 2.**
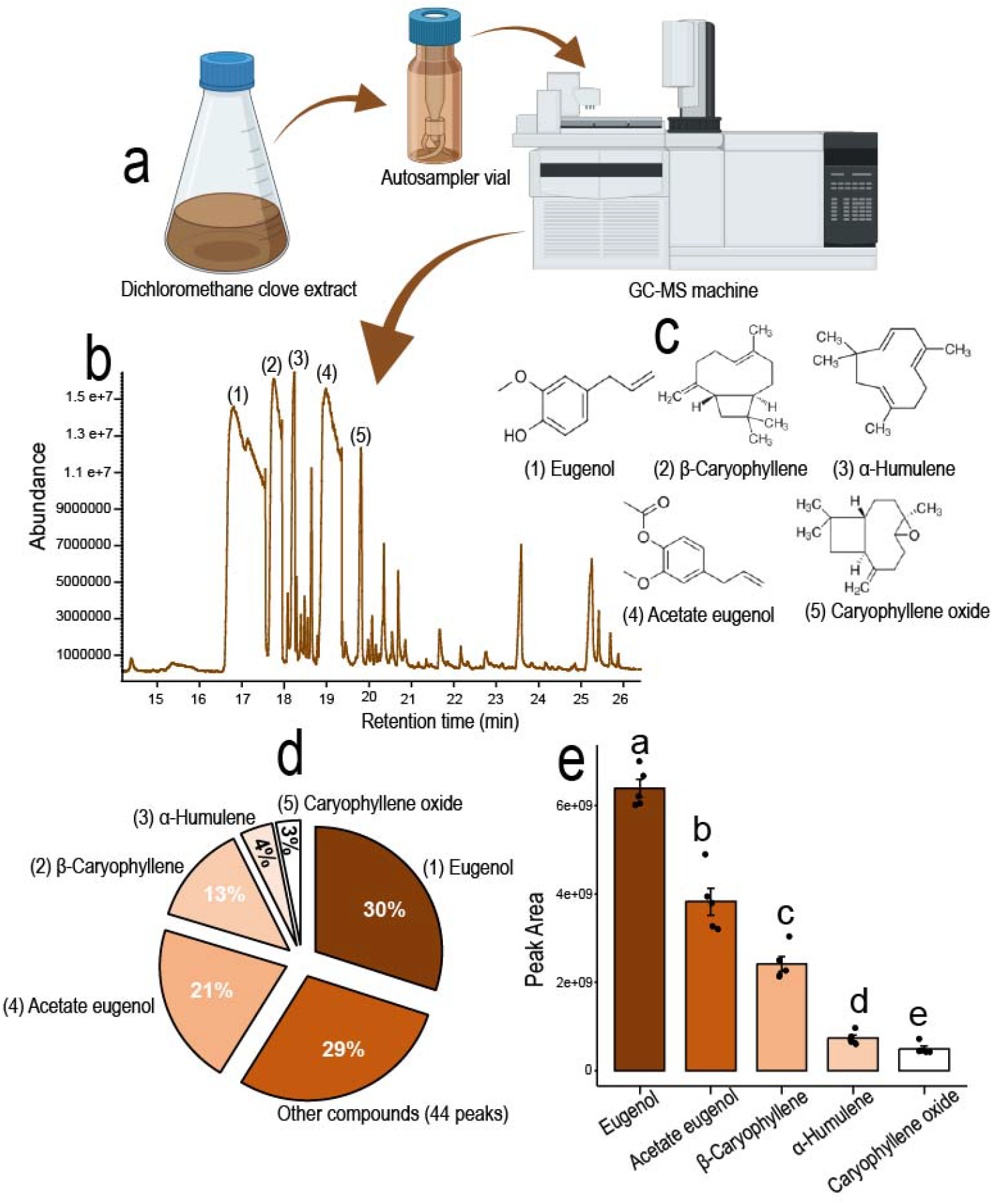
Eugenol is the major volatile in clove extract. **(a)** Extraction of volatiles from clove powder using dichloromethane. **(b)** Representative GC–MS chromatogram of clove volatiles. **(c)** Two-dimensional structures of the major volatile compounds identified from the clove extract. **(d)** Pie chart showing that eugenol represents the largest proportion among all detected volatiles. **(e)** Bar graph showing that eugenol had the highest mean peak area among the five major volatiles in the clove extract. Error bars indicate the standard error of the mean (SEM). Bars labelled with different letters differ significantly from each other.

### Behavioural Assays with synthetic volatiles

The chemical characterisation of clove extracts identified eugenol as the most abundant volatile compound. We therefore evaluated *D. suzukii’*s behavioural responses to synthetic eugenol using two-choice trap assays. Eugenol was diluted in mineral oil to various concentrations: undiluted, 10^− 1^, 10^− 2^, 10^− 3^, and 10^−4^ (w/v). For each assay, 200□µL of solution was placed in a cut Eppendorf tube lid positioned next to a half grape inside a 15□mL Nalgene specimen vial with 5□mm entry holes. The control consisted of a half grape near a cut Eppendorf lid containing 200□µL of mineral oil. Groups of 30 mixed-sex adult flies (3–5□days old, starved for 24□h) were gently released into cages measuring 15 cm × 15 cm × 20□cm, each containing the treatment and control (Fig. 3a). Trap positions were randomised to prevent side bias. After 24□h, the number of flies in each trap was counted. Each eugenol concentration was tested with 10 independent replicates using fresh flies and fruit. Repellency was assessed using a preference index (PI = (T − C)/(T + C)), where T and C represent flies on the treatment and control halves, respectively. Assays were conducted under controlled conditions (25 ± 1□°C, 65 ± 5% RH, 12:12□h light/dark cycle).

**Figure 3.**
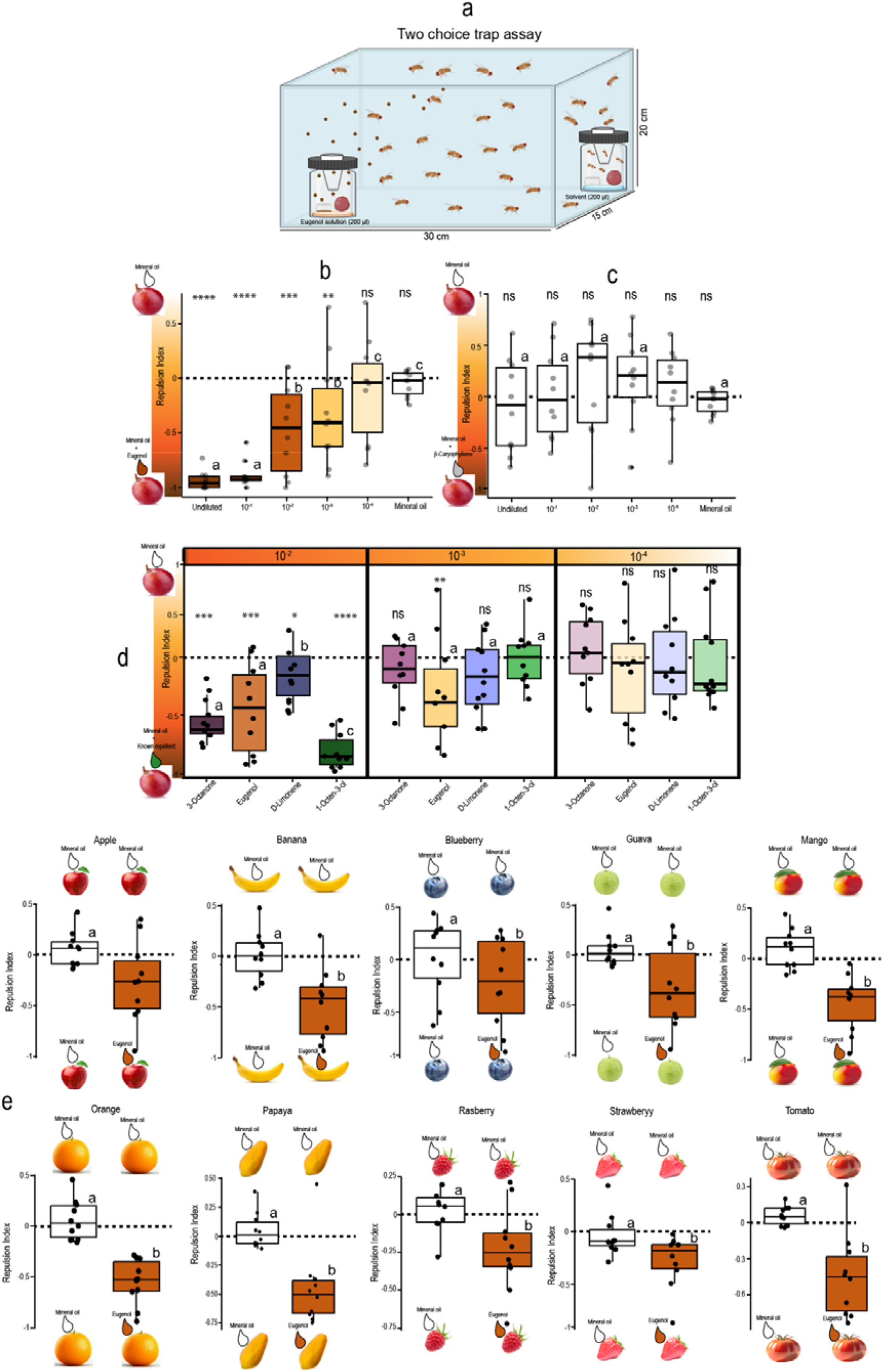
Eugenol elicits consistent avoidance behaviour in *Drosophila suzukii*. **(a)** Two-choice trap assay design. **(b)** Boxplots showing increased repellency of *D. suzukii* to grapes treated with eugenol at rising concentrations. **(c)** Boxplots showing no repellency of *D. suzukii* to grapes treated with (E)-β-caryophyllene across concentrations. **(d)** Boxplots comparing repellency of eugenol, 3-octanone, D-limonene, and 1-octen-3-ol at three concentrations (10^− 2^, 10^− 3^, 10^−4^). **(e)** Boxplots showing eugenol repellency across ten fruit hosts (apple, banana, blueberry, guava, mango, orange, papaya, raspberry, strawberry, and tomato). Boxplots show medians, interquartile ranges, and individual data points. “n.s.” indicates non-significant differences from the theoretical mean (0); *, **, ***, and **** indicate P□<□0.05, P□<□0.01, P□<□0.001, and P□<□0.0001, respectively. Different letters denote significant differences among treatments.

To contextualise eugenol’s efficacy, we also tested (E)-β-caryophyllene (>95% purity), a minor volatile of clove extract, at undiluted, 10^− 1^, 10^− 2^, 10^− 3^, and 10^−4^(w/v) concentrations. Additionally, we compared eugenol to previously identified *D. suzukii* repellents, including 3-octanone, 1-octen-3-ol (Wallingford et al., 2015; Xue et al., 2025), and (R)-(+)-limonene (Wang et al., 2021) at 10^− 2^, 10^− 3^, and 10^−4^. This comparison allowed us to benchmark eugenol’s repellent activity, validate assay sensitivity, and identify the most promising candidate for pest management.

Since fruits emit diverse volatiles that can synergise with or reduce repellency, we assessed eugenol’s effect across multiple host fruits, including apple, banana, blueberry, guava, orange, papaya, raspberry, strawberry, and tomato, using the same trap assay protocol. This ensured that eugenol’s repellency was consistent, ecologically relevant, and broadly applicable beyond grapes.

Highest-purity authentic standards of eugenol (≥98%), (E)-β-caryophyllene (>95%), 3-octanone (≥98%), 1-octen-3-ol (≥98%), (R)-(+)-limonene (≥98%), and mineral oil (≥99%) were purchased from Sigma-Aldrich (www.sigmaaldrich.com).

### Molecular Docking Procedure

Olfactory receptors are the primary proteins mediating odour detection in insects, making them the most relevant chemosensory targets (Fleischer et al., 2018; Wicher and Miazzi, 2021; Benton, 2022). Here, we aimed to elucidate the binding spectrum of eugenol to *D. suzukii* olfactory receptors. The complete chemosensory receptor repertoires in the *D. suzukii* genome have been characterised by Crava et al. (2016), Hickner et al. (2016), and Ramasamy et al. (2016). Thus, we obtained the nucleotide sequences of *D. suzikii* ORs from these studies and translated them into amino acid sequences using NCBI’s ORF Finder (https://www.ncbi.nlm.nih.gov/orffinder/). The 3D protein structures of these ORs were predicted with AlphaFold 2.0 (Jumper et al., 2021; Varadi et al., 2022). Protein structures were prepared for molecular docking in AutoDock Tools (Zhang et al., 2019) by removing water molecules, adding polar hydrogens, assigning Kollman charges, and applying Gasteiger charges. The 3D atomic structure of eugenol (SDF format) was retrieved from PubChem (https://pubchem.ncbi.nlm.nih.gov/compound/3314), optimised in Avogadro (Hanwell et al., 2012), and saved as a MOL2 file for docking. DeepSite (Jiménez et al., 2017) was used to predict potential binding sites on the OR structures, and the pocket coordinates were applied for docking in SwissDock (Bugnon et al., 2024), a web-based platform running AutoDock Vina (https://www.swissdock.ch/). Eugenol was docked against the complete *D. suzukii* OR repertoire. Binding affinities (kcal/mol) were recorded, and receptor– ligand interactions were ranked. In docking studies, binding free energy is a key measure of protein–ligand interactions, with more negative values indicating stronger binding and greater complex stability (Uzzaman & Uddin, 2019). This criterion was used to identify the ORs with the highest likelihood of strong eugenol binding. Finally, ChimeraX 1.10 (Meng et al., 2023) was used to visualise the binding modes and sites of eugenol within the most potent ORs.

### Electrophysiology

Our docking analysis revealed high binding affinity and stable interactions of eugenol with DsuzOR88a, DsuzOR69a, and DsuzOR23a. These receptors are expressed in olfactory sensory neurons housed in sensilla Dsuzat4, Dsuzab9, and Dsuzai2, respectively (Keesey et al., 2022). We therefore hypothesised that the predicted strong binding of eugenol to these proteins would translate into detectable physiological responses in vivo.

To test this, we performed single sensillum recordings (SSR) following the protocol of Baleba et al. (2023). Adult *D. suzukii* flies (2–4 days old) were immobilised in 100 µl plastic pipette tips, with only the head and antennae protruding. The preparation was secured on a microscope slide with dental wax, and the antenna was stabilised with a sharpened glass capillary positioned between the second and third antennal segments. Mounted flies were placed under an Olympus BX51WI light microscope equipped with a ×50 objective (LMPLFLN 50X) and 4× eyepieces. During recordings, the preparation was continuously flushed with charcoal-filtered, humidified air (1.5 l□min^−1^) delivered through a 0.4□cm tube positioned ∼6□cm from the fly.

Odour stimuli were delivered using cartridges prepared by loading a circular filter paper (1.2□cm diameter) with 10 µl of test solution and inserting it into a Pasteur pipette sealed with wax. Odour pulses (500□ms, 0.6□l□min ^−1^) were injected into the permanent airflow. Neural activity was recorded using electrochemically sharpened tungsten electrodes. The reference electrode was inserted into the eye, while the recording electrode was positioned at the base of a sensillum using a motor-controlled DC-3K micromanipulator (Märzhäuser) with a PM-10 piezo translator. Signals were amplified (USB-IDAC, Syntech) and analysed with AutoSpike software (version 3.7).

Guided by the sensilla distribution map of Keesey et al. (2022), we located the three target sensilla. To confirm their identity, we first stimulated them (10^-3^) with their strongest known ligands—methyl laurate (Dsuzat4C), linalool (Dsuzab9A), and farnesol (Dsuzai2B)—and observed robust excitatory responses, confirming correct placement. We then presented a solvent control (mineral oil) followed by a concentration series of eugenol (10^−5^ to 10^−1^). For each concentration of eugenol, recordings were made from one sensillum per fly, using a total of five flies. This experimental setup allowed us to directly test whether the strong binding predicted in silico would indeed translate into functional neuronal activation by eugenol.

### Cheminformatic Screening

We retrieved 1000 compounds structurally similar to eugenol (PubChem CID: 3314) from PubChem (accessed on 15 September 2025). SMILES representations of these molecules were used to calculate over 250 molecular descriptors with the rcdk R package (Guha, 2007), covering constitutional, topological, geometrical, electronic, and hybrid categories. Descriptors with near-zero variance were excluded, and highly correlated variables (Pearson’s r > 0.9) were removed to reduce redundancy. The remaining descriptors were standardised using a z-score transformation before downstream analyses. Cosine similarity indices (López-Pérez et al., 2024) were then computed to compare candidate molecules to eugenol, and the 10 most similar compounds were selected for docking experiments. These top candidates were docked against OR88a using the same pipeline as described above, and their binding affinities were compared with that of eugenol.

### Data analysis

All analyses were performed in R (version 4.5.1; R Core Team, 2025) using the RStudio interface (version 1.1.383). Graphs were assembled in Adobe Illustrator CC 2017 (version 21.0). Preference index data from the two-choice assays were analysed with a one-sample *t*-test when normally distributed (Shapiro–Wilk test: *P* > 0.05), or with a one-sample Wilcoxon test when not normally distributed (Shapiro–Wilk test: *P* < 0.05), comparing values against the theoretical mean of zero. Egg counts from the no-choice and multiple-choice oviposition assays were not normally distributed (Shapiro–Wilk test: *P* < 0.05) and their variances were not homogeneous (Barlet test: *P*<0.05). Therefore, we applied a Kruskal–Wallis test followed by Dunn’s test for pairwise comparisons (Dinno, 2024). Developmental time and pupal weight data were normally distributed with homogeneous (Bartlett’s test: *P*>0.05) variances. We analysed these data with ANOVA using the *agricolae* package (de Mendiburu, 2023), followed by the Student–Newman– Keuls (SNK) post hoc test to compare means across clove concentrations. Emergence percentage was analysed with a generalised linear model (GLM) using a binomial distribution (emerged vs. non-emerged) (Warton & Hui, 2011). Significance was assessed with analysis of deviance (chi-squared test), and pairwise differences among treatments were identified using Tukey’s test in the *emmeans* package (Lenth, 2025). Adult survival data were analysed with Kaplan–Meier survival analysis using the *survival* package (Therneau, 2024) and the *survminer* package (Kassambara et al., 2025), employing the functions survfit(), survdiff(), and pairwise_survdiff(). The relative proportions of clove volatile constituents were compared using chi-squared tests, while peak areas were compared with ANOVA followed by SNK post hoc tests. For the chemoinformatic analyses, we computed principal component analysis (PCA) on z-score–normalised molecular descriptor data from 1000 eugenol-like molecules. PCA was conducted with *FactoMineR* (Lê et al., 2008) and *factoextra* (Kassambara & Mundt, 2020), and the two-dimensional spatial distribution was visualised with *ggplot2* (Wickham, 2016).

## Results

### Clove extracts deter oviposition and reduce offspring performance in *D. suzukii*

We first tested the avoidance effect of clove on *D. suzukii* using a two-choice experiment. Results showed that flies strongly preferred control grapes over those treated with higher concentrations (10^− 1^, 10^− 2^, 10^− 3^ w/v), as reflected by negative preference indices (Fig. 1b). Gravid females showed the same avoidance during oviposition: they laid far fewer eggs on grapes coated with 10^− 1^ and 10^− 2^ clove solutions than on untreated grapes or those treated with the lowest concentration (10^−4^) (Fig. 1c). This pattern held in both no-choice assays (χ^2^ = 12.30, df = 4, P = 0.015; Fig. 1f) and multiple-choice assays (χ^2^ = 27.53, df = 4, P < 0.001; Fig. 1d), where egg deposition clearly declined with increasing clove concentration (Fig. 1d and e).

However, when females did oviposit on treated grapes, their offspring faced significant developmental costs. Larvae exposed to grapes coated with higher clove solution concentrations (10^− 1^ and 10^− 2^) developed more slowly than those on control grapes or lower dilutions (ANOVA, F_4,145_ = 8.13, P < 0.0001; Fig. 1f). Emerging adults were lighter in weight (ANOVA, F_4,81_ = 4.80, P = 0.011; Fig. 1g) and fewer in number (GLM, χ^2^ = 98.5, df = 4, P < 0.001; Fig. 1h), especially at 10^− 1^ and 10^− 2^ concentrations. Survival analyses confirmed these fitness costs: adults from clove-treated grapes had reduced lifespans compared to those from control grapes (log-rank test, χ^2^ = 13.24, P < 0.001; Fig. 1i).

Together, these results show that clove extracts affect *D. suzukii* at multiple stages. Adults avoid treated substrates, and when avoidance fails, their offspring experience slower growth, lower emergence, reduced body size, and shorter lifespans. Given these strong behavioural and fitness effects, we next asked which chemical components of clove extracts underlie this aversion.

### Chemical composition of clove volatiles

To better understand the chemical basis of the avoidance and life-history effects observed in *D. suzukii*, we analysed the volatile composition of the clove extract. Chemical profiling of the clove extract revealed a blend dominated by a handful of key volatiles. Gas chromatography analysis identified five major compounds: eugenol, (E)-β-caryophyllene, α-humulene, eugenol acetate, and caryophyllene oxide (Fig. 2b,c). Among them, eugenol stood out as the most abundant constituent, accounting for the largest proportion of the extract (χ^2^ = 42.56, df = 5, P = 0.01, Fig. 2d) and follow by eugenol acetate. Its dominance was further confirmed by quantitative comparisons of peak areas, where eugenol levels were significantly higher than those of the other compounds (ANOVA, F_4,20_ = 181.1, P < 0.001, Fig. 2e). Given that eugenol’s dominance, we next tested whether it drives the observed avoidance behaviour in the *D. suzukii*.

### Eugenol as the key clove-derived repellent in *D. suzukii*

We hypothesised that eugenol could be the main driver of the strong avoidance behaviour observed in *D. suzukii* toward clove-treated grapes. In two-choice trap assays, flies consistently avoided grapes placed near eugenol solutions (ANOVA, F_5,54_ = 12.78, P < 0.0001), with clear repellency at undiluted, 10^− 1^, 10^− 2^, and 10^− 3^ concentrations (Fig. 3b). To test whether this effect was specific, we next examined (E)-β-caryophyllene, another abundant constituent of clove extract after eugenol and its derivative acetate eugenol. Strikingly, flies showed no significant preference (ANOVA, F_5,54_ = 2.012, P = 0.0914), with distributions between treatment and control traps remaining close to equal (preference index ≈ 0) (Fig. 3c). This sharp contrast reinforced eugenol as the likely contributor to the aversive activity of clove extracts.

To place eugenol’s repellency into context, we compared its effects with three well-known insect repellents—3-octanone, D-limonene, and 1-octen-3-ol—across three concentrations (10^− 2^, 10^− 3^, and 10^−4^; Fig. 3d). At 10^− 2^ concentration, all four compounds reduced fly attraction, but their strength differed significantly (ANOVA, F_3,36_ = 10.08, P < 0.001). Limonene had the weakest effect, while eugenol, 3-octanone, and especially 1-octen-3-ol showed stronger repellency (Fig. 3d). At 10^− 3^ concentration, repellency did not differ significantly among compounds (ANOVA, F_3,36_ = 0.33, P = 0.803). However, one-sample t-tests revealed that only eugenol retained an effect significantly different from neutrality (t = -3.94, P < 0.05). At 10^−4^ concentration, none of the compounds differed significantly from neutrality (ANOVA, F_3,36_ = 0.33, P = 0.803). These findings highlight eugenol’s strong repellency at intermediate concentrations, whereas other compounds lose consistency.

Finally, to assess the ecological robustness of this response, we tested eugenol across a diverse panel of host fruits—apple, banana, blueberry, guava, mango, orange, papaya, raspberry, strawberry, and tomato. In every case, flies avoided eugenol-treated fruits, demonstrating a conserved aversion to this compound across substrates (Fig. 3e).

Together, these experiments reveal eugenol as the principal clove-derived volatile driving *D. suzukii* avoidance, with broad, consistent, and dose-dependent repellency across host environments.

### Molecular and physiological mechanisms of eugenol detection in *D. suzukii*

Olfactory receptors (ORs) are the molecular gatekeepers of odour detection, binding volatile molecules and converting them into neuronal signals that shape insect behaviour. To explore how *D. suzukii* perceives eugenol, we analysed its interactions with the predicted 3D structures of all reported *D. suzukii* Ors. Docking analyses highlighted three receptors with particularly relatively strong binding affinities: DsuzOR88a (-6.472kcal/mol), DsuzOR69a (–6.25 kcal/mol), and DsuzOR23a (–6.14 kcal/mol) (Fig. 4a) as compared to the binding affinity of the other receptors

**Figure 4.**
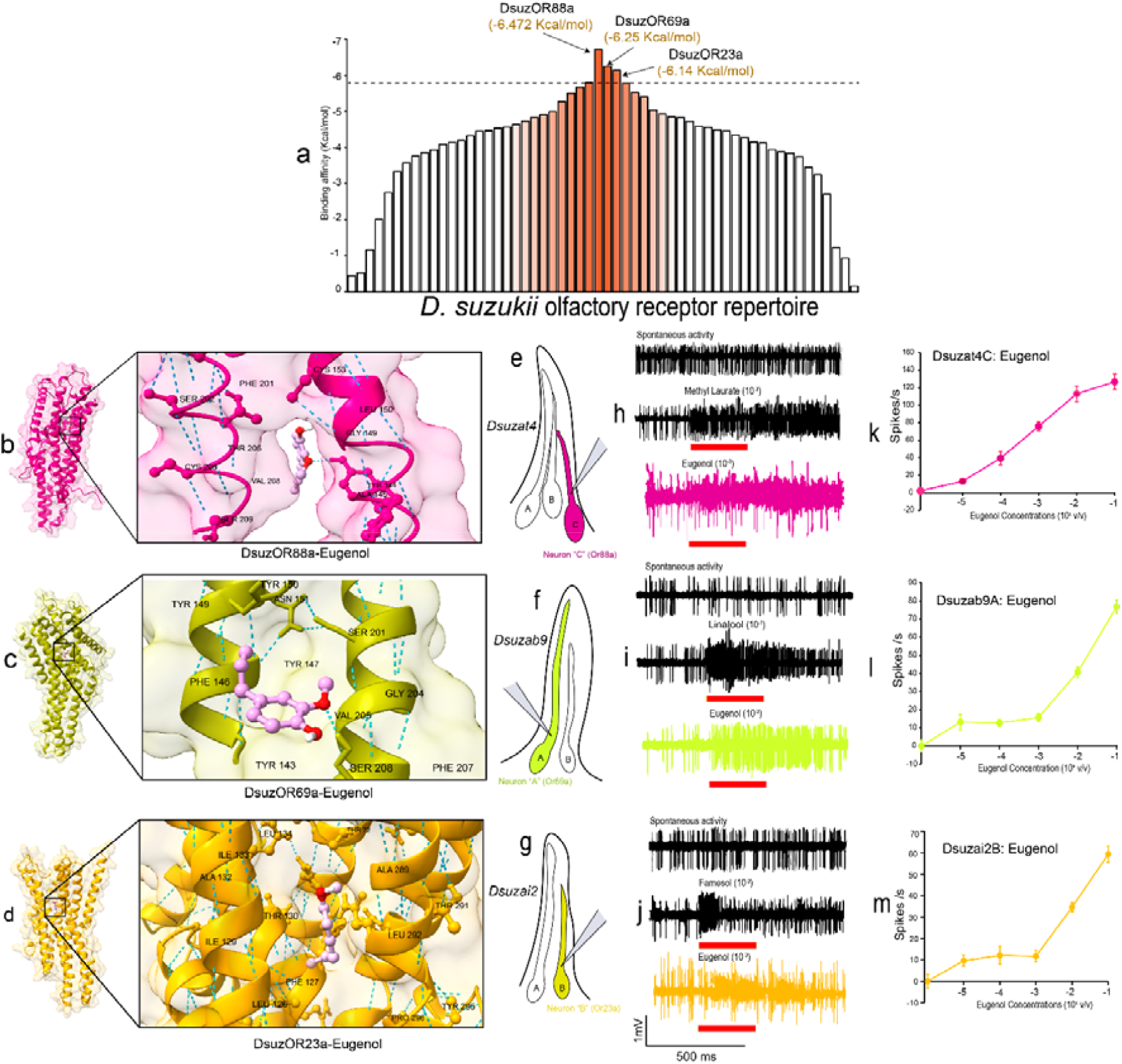
Molecular and physiological mechanisms of eugenol detection in *Drosophila suzukii*. **(a)** Tuning histogram showing docking affinities of eugenol with all predicted olfactory receptors. **(b–d)** Binding pocket architectures of DsuzOR88a, DsuzOR69a, and DsuzOR23a, highlighting key amino acid residues involved in eugenol stabilisation. **(e–g)** Sketches showing neurons C, A, and B expressing DsuzOR88a, DsuzOR69a, and DsuzOR23a in the at4, ab9, and ai2 sensilla, respectively. **(h–j)** Representative spike traces from single-sensillum recordings showing spontaneous neuronal activity, responses to diagnostic odours (10^− 3^), and responses to eugenol (10^− 3^) stimulation. **(k–m)** Line graphs showing dose-dependent responses of neurons C, A, and B to eugenol stimulation.

Each receptor presented a distinct binding environment. In DsuzOR88a (Fig. 4b), the pocket combined polar residues (SER209, THR205, SER202) with hydrophobic ones (VAL208, LEU150), supported by aromatic side chains (PHE201, TYR144). This composition created a balanced site where hydrogen bonds and π–π interactions stabilised eugenol binding. DsuzOR69a (Fig. 4c) displayed a contrasting architecture, dominated by aromatic residues, including multiple tyrosines (TYR143, TYR147, TYR149, TYR150) and phenylalanines (PHE146, PHE207). This aromatic-rich pocket was well suited for π–π stacking with eugenol’s phenolic ring, while polar residues (ASN151, SER201) contributed to hydrogen-bonding contacts, explaining the observed strong affinity. In contrast, DsuzOR23a (Fig. 4d) exhibited a largely hydrophobic architecture, lined with leucines (LEU126, LEU134, LEU292), isoleucines (ILE129, ILE133), and alanines (ALA132, ALA289). This nonpolar environment accommodated the hydrocarbon tail of eugenol, while TYR295 and threonines (THR130, THR291) introduced specific polar contacts that fine-tuned the interaction. Together, these results demonstrate how *D. suzukii* ORs, despite structural differences, converge on eugenol recognition through unique combinations of hydrophobic, polar, and aromatic residues, highlighting the molecular versatility of the insect olfactory system.

Having established strong ligand–receptor affinities, we next tested whether these interactions produced physiological activity. In *D. suzukii*, DsuzOR88a is expressed in the C neuron of at4 sensilla (Fig. 4e), DsuzOR69a in the A neuron of ab9 sensilla (Fig. 4f), and DsuzOR23a in the B neuron of ai2 sensilla (Fig. 4g). Using the sensilla distribution map of Keesey et al. (2022), we located these structures on the antenna and verified their identity electrophysiologically. Diagnostic ligands methyl laurate for at4 (Fig. 4h), linalool for ab9 (Fig. 4i), and farnesol for ai2 (Fig. 4j)—elicited characteristic spike patterns in the expected neurons.

Subsequent stimulation with eugenol induced clear, dose-dependent increases in action potentials across all three neurons: C (at4), A (ab9), and B (ai2) (Figs. 4k–m). Notably, neuron C produced the highest spike frequency, followed by A and B, mirroring the receptor affinity pattern from docking analyses. This congruence between in silico and in vivo results strengthens the causal link between molecular binding and neuronal activation.

Altogether, these findings reveal that eugenol detection in *D. suzukii* is mediated by a small set of highly responsive ORs whose ligand–receptor interactions drive robust neural responses. This mechanistic insight bridges molecular recognition with behavioural avoidance, explaining how a single plant-derived compound can trigger strong repellent behaviour through precise olfactory coding.

### Identification of eugenol-like repellents through chemoinformatic screening

Having demonstrated that eugenol activates specific olfactory receptors in *D. suzukii*— particularly DsuzOR88a, followed by DsuzOR69a, and DsuzOR23a—through distinct molecular interactions resulting in strong neuronal activity, we then investigated whether structurally related molecules could induce similar effects. To do this, we carried out a large-scale chemoinformatic screen of 1000 eugenol-like molecules downloaded from PubChem (Fig. **5a**). Each molecule was characterised by molecular descriptors (Fig. **5b**), and their similarity to eugenol was measured using the cosine similarity index (Fig. **5c**).

**Figure 5.**
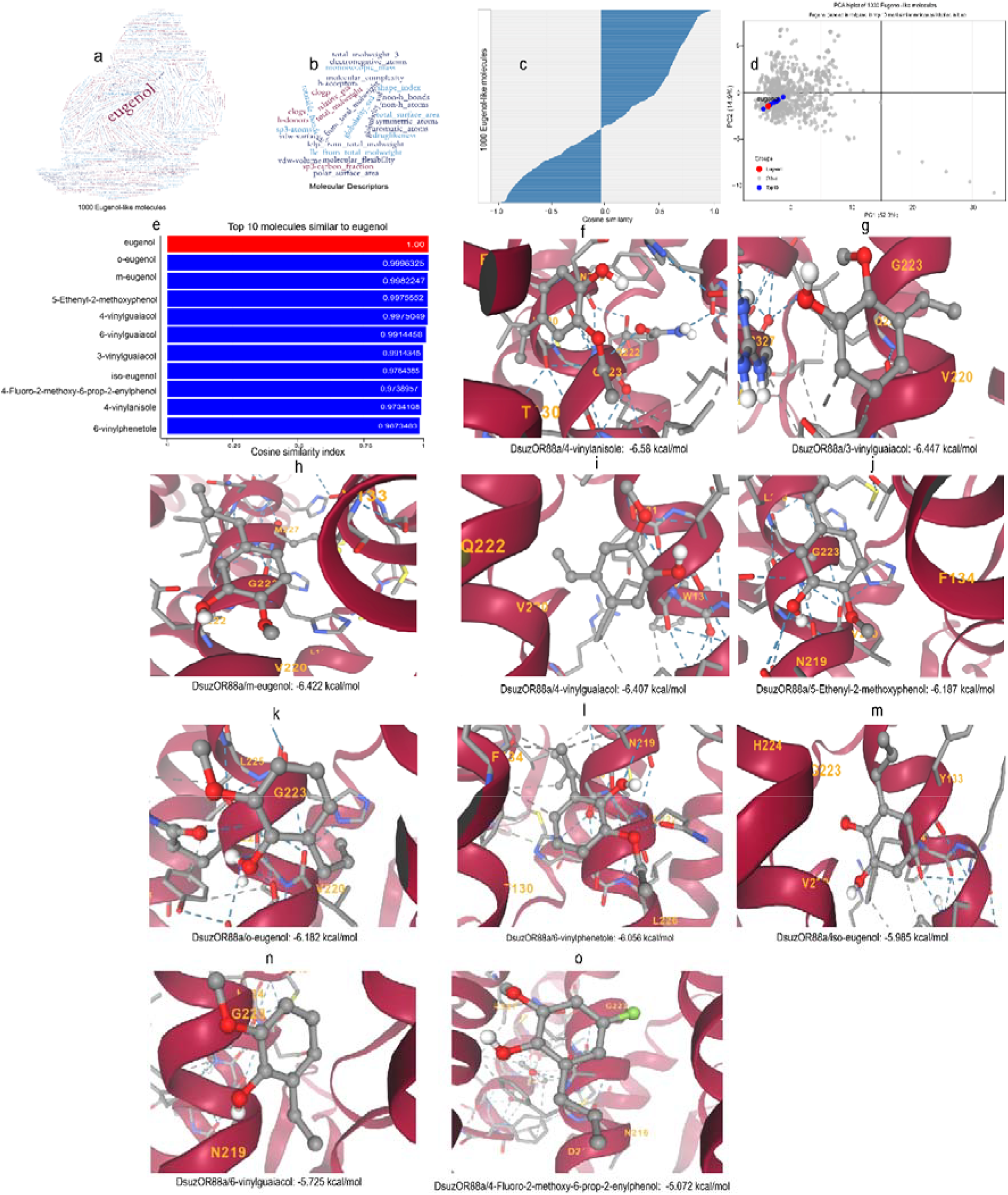
Chemoinformatic and molecular docking analysis of eugenol-like molecules in *Drosophila suzukii* OR88a. **(a)** Word cloud of 1000 eugenol-like molecules retrieved from PubChem. **(b)** Word cloud of molecular descriptors computed for the 1000 compounds. **(c)** Distribution of cosine similarity indices showing structural similarity of each compound to eugenol. **(d)** Principal component analysis (PCA) of the 1000 compounds based on molecular descriptors, highlighting clustering of eugenol with its closest analogues. **(e)** Histogram ranking the ten most structurally similar compounds to eugenol based on their cosine similarity indices. **(f–o)** Molecular docking poses of the top ten eugenol analogues within the DsuzOR88a binding pocket, showing key interactions and predicted binding affinities (kcal/mol).

Principal component analysis (PCA) offered a two-dimensional view of the chemical space, with the first two axes accounting for 52.3% (PC1) and 14.9% (PC2) of the variance (Fig. **5d**). Within this space, eugenol clustered closely with a group of structurally similar compounds. The top ten most similar compounds to eugenol were *o*-eugenol (cosine index: 0.9996), *m*-eugenol (0.9982), 5-ethenyl-2-methoxyphenol (0.9976), 4-vinylguaiacol (0.9975), 6-vinylguaiacol (0.9914), 3-vinylguaiacol (0.9914), iso-eugenol (0.9784), 4-fluoro-2-methoxy-6-prop-2-enylphenol (0.9739), 4-vinylanisole (0.9734), and 6-vinylphenetole (0.9673) (Fig. **5e**)

Encouraged by these results, we examined whether the selected eugenol analogues interact with *Drosophila suzukii*’s olfactory receptor DsuzOR88a, one of eugenol’s primary targets. Docking simulations revealed that 4-vinylanisole (–6.580 kcal/mol, Fig. **5f**) exhibited the strongest binding among the tested analogues, surpassing eugenol (–6.472 kcal/mol). This was followed by 3-vinylguaiacol (–6.447 kcal/mol, Fig. **5g**), *m*-eugenol (–6.422 kcal/mol, Fig. **5h**), and 4-vinylguaiacol (–6.409 kcal/mol, Fig. **5i**), all showing comparable binding strengths to eugenol. Compounds with slightly weaker affinities included 5-ethenyl-2-methoxyphenol (–6.187 kcal/mol, Fig. **5j**), *o*-eugenol (–6.182 kcal/mol, Fig. **5k**), and 6-vinylphenetole (–6.056 kcal/mol, Fig. **5l**). The weakest interactions were observed for iso-eugenol (–5.985 kcal/mol, Fig. **5m**), 6-vinylguaiacol (–5.725 kcal/mol, Fig. **5n**), and 4-fluoro-2-methoxy-6-prop-2-enylphenol (–5.097 kcal/mol, Fig. **5o**). These results suggest that variations in the position or type of substituent groups on the aromatic ring influence receptor binding strength, highlighting the importance of hydroxyl and methoxy group orientation in determining ligand–receptor affinity.

Together, these results show that several eugenol analogues not only share high structural similarity with eugenol but also bind DsuzOR88a with equal or stronger affinity. This suggests functional overlap or diversification in olfactory coding. The combined chemical similarity and receptor docking analyses highlight a family of eugenol-like molecules with potential as novel repellents, offering new candidates for disrupting *D. suzukii* host-finding behaviour. These findings identify a family of eugenol-like molecules as promising candidates for developing scent-based strategies to disrupt *D. suzukii* host-finding, with potential applications in sustainable pest management.

## Discussion

*Drosophila suzukii* presents a distinct threat to fruit production because, unlike most Drosophila species that target decaying fruit, it oviposits in healthy, ripening fruit. This behaviour results in significant yield losses, diminished market value, and increased management costs (Bolda et al., 2010; Walsh et al., 2011; Asplen et al., 2015). As reliance on prophylactic insecticide sprays proves unsustainable, innovative strategies are urgently required. Our findings show that clove volatiles, particularly eugenol, repel adult *D. suzukii* while also inhibiting the development of their offspring, suggesting a dual-action approach with strong potential for integrated pest management (IPM). To our knowledge, no previous study has reported that clove extracts or eugenol simultaneously deter *D. suzukii* adults and impair offspring development, underscoring the novelty and significance of these results.

Our results show that clove extracts deter *D. suzukii* at multiple stages. Adults avoided grapes treated with moderate to high concentrations, and gravid females laid fewer eggs on treated fruits. Offspring developing on treated grapes exhibited slower growth, reduced emergence, smaller body size, and shorter lifespans. This dual action indicates that clove compounds influence both behaviour and fitness. Previous studies with other plant volatiles reported mainly behavioural effects. Peppermint oil reduced adult attraction (Renkema et al., 2016), while Lavandula latifolia and avocado extracts deterred oviposition and reduced survival (Erland et al., 2015). Similarly, Souza et al. (2022) found that 90% of tested plant extracts repelled *D. suzukii* females, with *Cymbopogon flexuosus* and *Mentha* spp. extracts showing strong toxicity. Unlike these studies, our findings highlight direct impacts on larval development and adult fitness. Comparable effects have been reported in other insects: neem extracts reduced growth and fecundity in *Spodoptera frugiperda* (Martinez & van Emden, 2001), and garlic oil reduced survival *in Tenebrio molitor* larvae (Plata-Rueda et al., 2017). Evidence for such developmental penalties in *D. suzukii* remains scarce, making clove extracts a novel candidate for managing this fly. These effects were dose-dependent, with stronger avoidance and developmental costs at higher concentrations. Similar dose–response patterns have been observed for peppermint and geranium oils (Renkema et al., 2016), emphasising the importance of optimising field application rates. Overall, these results demonstrate that clove extracts offer dual benefits—reducing oviposition and impairing offspring performance—thereby increasing their potential utility in integrated pest management strategies. This combination of behavioural repellency and developmental suppression represents the first documented dual-action effect of a plant-derived compound against *D. suzukii*, providing a new avenue for sustainable control.

Within clove volatiles, chemical profiling identified eugenol as the dominant compound, consistent with previous reports. For example, Haro-González et al. (2021) showed that eugenol accounts for at least 50% of clove oil, with eugenyl acetate, β-caryophyllene, and α-humulene making up the remaining 10–40%. Eugenol is a phenylpropanoid with diverse biological activities, including insecticidal, antimicrobial, anti-inflammatory, antiviral, and antioxidant effects (Ulanowska & Olas, 2021; Silva et al., 2024). Beyond these known properties, our experiments revealed that eugenol is a potent repellent against *D. suzukii*. Strikingly, this repellency was not limited to grapes: when tested across a panel of ten host fruits, including apple, banana, blueberry, guava, mango, orange, papaya, raspberry, strawberry, tomato, eugenol consistently repelled *D. suzukii*. Such consistency suggests evolutionary pressure to maintain sensitivity to this compound across ecological contexts. This conserved aversion across substrates demonstrates the ecological robustness of the response and strengthens the case for eugenol as a broadly effective repellent under field conditions. Additionally, this trend could reflect co-evolutionary interactions where host fruits bearing eugenol-related volatiles are non-preferred and therefore lead to the conservation of avoidance circuits in *D. suzukii* odorant pathways. The broad-spectrum repellency would then be an adaptive sensory bias on the basis of the perception of plant secondary metabolites. The demonstration that eugenol repels *D. suzukii* across ten fruit types is unprecedented, indicating a broad-spectrum ecological robustness not previously described for any single plant volatile.

Eugenol’s repellency has been observed across various insect species, including *Aedes albopictus* (Deng et al., 2023), *Sitophilus oryzae* (Abenaim et al., 2022), *Solenopsis invicta* (Wang et al., 2024), and *Musca domestica* (Subaharan et al., 2021). This repellency is closely linked to its insecticidal properties. For example, Obeng-Ofori and Reichmuth (2017) demonstrated the strong toxicity of eugenol when applied topically, impregnated on filter papers, or mixed with whole grains, affecting *Sitophilus granarius, S. zeamais, Tribolium castaneum*, and *Prostephanus truncatus*. In *M. domestica*, contact assays revealed that eugenol had high larvicidal potential, exerted pupicidal effects, and caused adult malformations when treated pupae. Consistent with these findings, we observed prolonged development, reduced emergence, smaller body size, and lower survival rates in *D. suzukii* offspring from grapes treated with clove extract. Since eugenol was identified as the dominant component in our chemical analysis, these adverse effects are most likely attributable to its action. Mechanistic studies further support multiple molecular targets for eugenol in insects. In *Triatoma infestans*, topical eugenol triggered hyperactivity that was mitigated by phentolamine, suggesting octopamine receptors as one site of action (Reynoso et al., 2020). In *Periplaneta americana* and *Drosophila melanogaster*, eugenol and related essential oil constituents disrupted binding to cloned octopamine receptors (Paoa 1, OAMB), altered intracellular cAMP and calcium signalling, and these receptor-mediated effects correlated with toxicity (Enan, 2005). Additionally, in *Psoroptes cuniculi* and insect-derived cell lines such as *Aedes albopictus* C 6/36, eugenol inhibited mitochondrial complex I (NADH dehydrogenase) activity and disrupted mitochondrial membrane potential, displaying selective toxicity towards insect and arthropod cells compared with mammalian cells (Shang et al., 2020). All of this makes eugenol a reliable chemical “danger cue”. By detecting and avoiding eugenol, *D. suzukii* reduces the risk of exposing itself and its offspring to toxic environments, as evidenced by our observation of decreased survival and fitness in larvae developing on eugenol-treated fruits. This could also highlight the evolutionary concept of semiochemicals as ‘eco-toxicological signals,’ where detection of a compound warns insects of metabolically incompatible environments. Collectively, these findings show that eugenol, the primary component of clove extract, exhibits broad-spectrum repellency and toxicity, acting through multiple molecular pathways. Its ability to repel *D. suzukii* across various host fruits while simultaneously impairing offspring performance highlights its ecological robustness and underlines its potential as a plant-derived tool for sustainable pest management. These findings constitute the first comprehensive evidence linking eugenol’s molecular targets with both behavioural and developmental outcomes in *D. suzukii*, extending its known mode of action beyond other insect taxa.

Olfaction is central to *D. suzukii* host location and oviposition (Keesey et al., 2015, 2022). Olfactory receptors (ORs) mediate this process by binding volatile molecules and converting them into neuronal signals that guide behaviour (Andersson et al., 2015). Our docking analyses showed that among all the *D. suzukii* olfactory receptors, eugenol interacts most strongly with DsuzOR88a (−6.86 kcal/mol), followed by DsuzOR69a (−6.25 kcal/mol) and DsuzOR23a (−6.14 kcal/mol). The binding pocket of DsuzOR88a combines polar (SER209, THR205, SER202), hydrophobic (VAL208, LEU150), and aromatic (PHE201, TYR144) residues, whereas DsuzOR69a is dominated by aromatics (TYR143, TYR147, TYR149, TYR150, PHE146, PHE207) with polar inputs (ASN151, SER201), and DsuzOR23a is largely hydrophobic (LEU126, LEU134, LEU292, ILE129, ILE133, ALA132, ALA289) with selective polar–aromatic contacts (TYR295, THR130, THR291). This explains why OR88a binds eugenol most strongly, as its balanced pocket supports both hydrogen bonding and π–π interactions, a mechanism also described for MhOR5 in the jumping bristletail *Machilis hrabei* (Del Mármol et al., 2021). Strong support for this interpretation comes from *Bactrocera dorsalis*, where Liu et al. (2018) identified BdorOR88a as the primary receptor for methyl eugenol, a structural analogue of eugenol. They showed that BdorOR88a elicited robust responses to methyl eugenol in Xenopus oocytes, and silencing this receptor significantly reduced male attraction to the compound. Similarly, in our single sensillum recording assays, stimulation of neuron “C” expressing DsuzOR88a produced action potentials in a dose-dependent manner. This conserved role of OR88a in detecting eugenol-related ligands across species supports the high binding affinity observed for DsuzOR88a and underscores its ecological importance in *D. suzukii* host-odour detection. Notably, BdorOR88a is an orthologue of DmelOR88a in *Drosophila melanogaster* (Liu et al., 2018), which is in turn orthologous to DsuzOR88a in *D. suzukii*, highlighting a conserved evolutionary role of OR88a in detecting eugenol-like compounds across dipteran species (Walker et al., 2023). This is the first functional and structural evidence implicating DsuzOR88a in eugenol detection, revealing a previously uncharacterised molecular mechanism of odour recognition in *D. suzukii*.

We also observed that neurons “A” and “B”, which express DsuzOR69a and DsuzOR23a, the second and third strongest eugenol-binding receptors, generated action potentials upon eugenol stimulation, though with differing spike counts. This detection of one specific odour by multiple ORs has been observed in several insect species, including *D. melanogaster* (Hallem and Carlson, 2006), *Harpegnathos saltator* (Slone et al., 2017), *Spodoptera littoralis* (De Fouchier et al., 2017), *Locusta migratoria* (Chang et al., 2023) and *Aedes aegypti* (Singh et al., 2023). Our findings thus support the combinatorial coding model of odour detection in insects as reviewed by Haverkamp et al. (2023).

In *D. suzukii*, neurons “C”, “A”, and “B” express DsuzOR88a, DsuzOR69a, and DsuzOR23a, and are housed in antennal sensilla trichoid type 4 (at4), basiconic type 9 (ab9), and intermediate type 2 sensilla (ai2), respectively (Keesey et al., 2022). Neuron C in at4 trichoid sensilla expresses DsuzOR88a, which showed the strongest binding affinity for eugenol and a clear electrophysiological response. This matches the established role of trichoid sensilla in detecting host-related volatiles and pheromone-like compounds. For illustrations, Pitts et al. (2016) demonstrated that Or88a neurons in *D. melanogaster* are sensitive to a diverse array of ligands, including both fly and non-fly derived odours. Neuron A in ab9 basiconic sensilla expresses DsuzOR69a, which displayed moderate binding affinity to eugenol. Basiconics typically detect plant volatiles and food odours, suggesting a secondary but ecologically relevant role for DsuzOR69a in mediating eugenol-driven host attraction. In support, the DoOR database (https://neuro.uni-konstanz.de/DoOR/default.html) that offers a comprehensive mapping of *D. melanogaster* odour response (Münch et al., 2016), lists eugenol among the best ligands of DmelOR69a, the orthologue of DsuzOR69a. Neuron B in intermediate type 2 sensilla expresses DsuzOR23a, the weakest binder, which may explain its reduced contribution to eugenol detection compared to OR88a and OR69a. By demonstrating that eugenol activates multiple olfactory receptor neurons in *D. suzukii*, our study provides new experimental support for combinatorial coding of plant volatiles in this pest species.

Together, these results suggest that eugenol detection in *D. suzukii* is primarily mediated by DsuzOR88a, supported by DsuzOR69a, while DsuzOR23a plays a minor role. This division of labour across sensilla types highlights how *D. suzukii* integrates signals from multiple sensory channels to fine-tune host-odour detection and oviposition behaviour. These receptor-level insights represent a new step in understanding odour-guided host selection in *D. suzukii*, linking molecular, physiological, and behavioural responses for the first time.

Having demonstrated that eugenol activates specific olfactory receptors in *Drosophila suzukii*, with DsuzOR88a showing the strongest neuronal response, we next examined whether structurally related molecules could produce similar effects. A large-scale chemoinformatic screening of 1000 eugenol-like molecules, based on molecular descriptors and cosine similarity index, identified the ten most similar compounds: *o*-eugenol, *m*-eugenol, 5-ethenyl-2-methoxyphenol, 4-vinylguaiacol, 6-vinylguaiacol, 3-vinylguaiacol, iso-eugenol, 4-fluoro-2-methoxy-6-prop-2-enylphenol, 4-vinylanisole, and 6-vinylphenetole. Most of these compounds share the key structural features responsible for receptor activation, a phenolic ring bearing a methoxy (-OCH_3_) group and an allyl or propenyl side chain, explaining their close similarity to eugenol in both structure and binding affinity. These moieties are known to influence both molecular binding affinity and receptor selectivity in insect olfactory systems (Hallem and Carlson, 2006; Pellegrino et al., 2011; Dweck et al., 2013). In support of this, our docking simulations revealed that several analogues bind DsuzOR88a with comparable affinity to eugenol (–-6.472 kcal/mol). Among these top ten analogues, 4-vinylanisole showed the highest predicted affinity (-6.58 kcal/mol), followed by 3-vinylguaiacol (–6.447 kcal/mol), m-eugenol (–-6.442 kcal/mol), and 4-vinylguaiacol (–6.409 kcal/mol). These results suggest that DsuzOR88a possesses a flexible binding pocket capable of accommodating minor structural variations while preserving receptor–ligand interactions. Such receptor promiscuity may allow *D. suzukii* to detect a broader range of host-related volatiles, reflecting an adaptive advantage in complex odour landscapes. This finding is consistent with studies showing that odorant receptors often recognise families of structurally related compounds rather than single ligands. This pattern has been well documented across insects and mammals. In *Drosophila*, Hallem and Carlson (2006) showed that many ORs respond to groups of odorants sharing core structural motifs, reflecting broad tuning within chemical families. Similar multi-ligand recognition has been described in *Anopheles gambiae* ORs, which detect related host-derived volatiles (Carey et al., 2010), and in mammalian receptors following a combinatorial coding scheme (Malnic et al., 1999). Comparative analyses and chemoinformatic studies further confirm that receptor activation is strongly predicted by molecular similarity among ligands (Bohbot & Dickens, 2012; Boyle et al., 2013). This study provides the first evidence that multiple structurally related compounds can target DsuzOR88a, a receptor identified as important for eugenol detection in *D. suzukii*. The demonstration that eugenol and its analogues share receptor-level recognition patterns expands the current understanding of odour detection mechanisms and supports the idea of combinatorial coding in insect olfaction. In addition to its biological significance, the integration of chemoinformatics with receptor docking represents a novel methodological approach for predicting odour-receptor interactions in non-model insect species. By combining molecular similarity screening and computational docking, this study establishes a predictive framework for identifying new semiochemicals with repellent potential prior to behavioural or electrophysiological testing. For illustration, 3-vinylguaiacol, 4-vinylguaiacol and 6-vinylguaiacol, which rank among the top ten eugenol-like molecules identified, are analogues to guaiacol, a molecule that repels three Drosophila species, namely *D. ezoana, D. novamexicana* and *D. virilis* (Baleba et al., 2023). Guaiacol also forms part of the formulation of Waterbuck Repellent Blend that repels tsetse flies (Bett et al., 2015; Diallo et al., 2020). Ecologically, identifying eugenol analogues that match eugenol’s affinity for DsuzOR88a opens new avenues for developing environmentally safe, scent-based repellents. The discovery of these analogues is therefore not only chemically and mechanistically novel but also highly relevant for integrated pest management (IPM). Collectively, the combined chemical similarity and receptor docking analyses highlight a family of eugenol-like molecules with potential as novel repellents. These findings identify new candidate compounds that could be exploited to design scent-based strategies to interfere with *D. suzukii* host location, providing a promising step toward sustainable pest control solutions.

## Conclusion

Our study identifies eugenol as a potent dual-action compound against *Drosophila suzukii*. It repels adults across fruit hosts and suppresses larval development, emergence, and survival. This dual activity positions clove-derived eugenol as a promising candidate for sustainable, plant-based pest management. By integrating chemical profiling, electrophysiology, molecular docking, and chemoinformatics, we show that eugenol detection is mainly mediated by DsuzOR88a, with DsuzOR69a and DsuzOR23a providing complementary detection through distinct sensilla types. Chemoinformatic screening further reveals that several eugenol analogues bind DsuzOR88a with comparable affinities, highlighting receptor flexibility and potential for designing new repellents based on structural similarity. This receptor-level insight provides a mechanistic basis for eugenol avoidance and supports the development of scent-based, environmentally safe pest control strategies. Eugenol and its analogues can disrupt host finding and reproduction through specific olfactory pathways, offering both molecular and ecological leverage. Future work should test eugenol (including its analogues we identified) and clove extracts under field conditions across different crops, climates, and seasons. Developing slow-release or micro-encapsulated formulations may enhance stability, while blending eugenol with other volatiles could increase efficacy. Structure–activity studies should identify analogues with stronger DsuzOR88a binding and improved selectivity. Further molecular validation, including receptor expression assays and CRISPR-Cas9 knockouts, can confirm the roles of DsuzOR88a, DsuzOR69a, and DsuzOR23a. Transcriptomic profiling could reveal downstream signalling pathways, and behavioural assays should test how eugenol affects oviposition, learning, and mating across sexes and species. Toxicological and compatibility tests will clarify eugenol’s safety for non-target insects and its integration with biological control agents. Evolutionary studies could also assess whether long-term exposure drives receptor adaptation and how eugenol functions as a defensive or warning cue in natural host–plant interactions. Our receptor models are computational predictions and may not fully capture native receptor dynamics. Docking scores approximate binding free energy but may not represent in vivo conditions. Future research should combine molecular dynamics, mutagenesis, and in vivo assays to validate ligand selectivity and refine the understanding of *D. suzukii* olfactory mechanisms.

## Author Contributions

**Steve B. S. Baleba:** Conceptualised the study, collected, curated, and analysed the data, prepared the figures, and wrote the first draft of the manuscript. **Victor O. Omondi:** Collected data on the chemical analysis of clove extracts and contributed to manuscript review and editing. **Samira A. Mohamed:** Provided access to the *Drosophila suzukii* colony and contributed to manuscript review and editing. **Merid N. Getahun:** Reviewed and edited the manuscript. **Nan-ji Jiang:** Collected data, prepared figures, and contributed to manuscript review and editing. **Souleymane Diallo:** Advised on the molecular docking protocol and contributed to manuscript review and editing.

## Acknowledgements

This project did not receive specific funding from any research grant. It was supported by the personal salary of Steve B. S. Baleba, paid through the core funding of icipe by its donors, including the Swedish International Development Cooperation Agency (Sida), the Swiss Agency for Development and Cooperation (SDC), the Australian Centre for International Agricultural Research (ACIAR), the Government of Norway, the German Federal Ministry for Economic Cooperation and Development (BMZ), and the Government of the Republic of Kenya. The authors acknowledge these donors for their support. We thank Ms Fridah M. Kasolo for providing the flies used in this study. The views expressed here do not necessarily reflect those of the donors.

